# A novel rat model of body first subtype of Parkinson’s disease

**DOI:** 10.64898/2026.07.13.738258

**Authors:** Vaibhavi Peshattiwar, Caroline Swain, Dipesh Pokharel, Khoi Le, Isabel Kennedy, Kala Venkiteswaran, Thyagarajan Subramanian

## Abstract

The recent growing evidence support the existence of two subtypes of Parkinson’s disease (PD), a body first and brain first subtype owing to variability in the disease site of onset as well as disease progression. Animal models which could replicate the specific differences of these subtypes are important to explore the pathophysiology as well as to evaluate novel treatment options. Here, we describe an animal model of body first PD subtype developed using repeated low dose exposure of environmental neurotoxin Paraquat (P) and Lectin (L) to characterize its PD-like manifestations. We administered P+L (P+L, p.o.) daily to rats for 90 days. These animals underwent motor and non-motor behavioral tests at various time intervals. After 21 weeks, post-mortem histopathological analysis was performed to assess neurodegeneration. Onset of motor deficits initiated unilaterally from week 4 of P+L followed by gradual progression towards bilateral symptoms that were levodopa responsive. This model also replicates non motor features including cognitive deficits in tests like Novel Object Recognition Test and Y maze as well as sleep abnormalities. The histopathology showed nigrostriatal dopaminergic degeneration and proteinase K resistant S129 alpha-synucleinopathy both in the gut and the brain. The replication of both progressive motor and non-motor features in this rat model corroborates body first subtype of PD therefore making it an attractive option for testing neuroprotective experimental therapeutics and avenue to understand pathophysiological mechanisms.

## Introduction

Parkinson’s disease (PD) have been established as an heterogenous, multifactorial progressive neurodegenerative disorder. Recent data from various postmortem and imaging studies which points towards the clinical heterogeneity has led researchers to differentiate PD into body and brain first subtypes. This differentiation assimilates the distinct phenotype as well as origins of α-synuclein pathology^1–5^. Though recent findings have led to evolved understanding of idiopathic PD, the exact etiology and pathophysiology is still elusive. Therefore, there is urgent need for further investigations directed to address many fundamental questions related to PD pathogenesis. Preclinical evaluation have played a critical role in unraveling pathophysiological mechanisms as well as translational therapeutic research aiding clinical investigations. A prevailing challenge currently in the preclinical translational research for idiopathic PD is development of an animal model which could faithfully replicate the subtype specific pathology and progression including the associated risk factors of PD. One of the important risk factors reported in these findings are the environmental neurotoxicants especially a widely used herbicide Paraquat. Several epidemiological studies have established a link between the development of idiopathic PD and Paraquat (P) ^6^.

We have shown previously that oral co-administration of subthreshold doses of P along with lectins (L), a ubiquitous carbohydrate-binding dietary protein (from Pisum sativum) in adult rats over a 7-day period causes bilateral parkinsonism that is responsive to L-dopa treatment and can be completely prevented by bilateral sub-diaphragmatic vagotomy ^7^ These animals exhibited slowing of gastric motility and pathological evidence of Serine129 phosphorylated alpha synucleinopathy in the myenteric plexus, the dorsal motor nucleus of the vagus and in the substantia nigra pars compacta in the brain with associated >45% loss of dopaminergic neurons and its projections^8^. To improve upon this model and to emulate a more likely scenario of human exposure we simulated the longer exposure time of 90 days at lower daily dose while maintaining the total cumulative dose the same as in the 7 days exposure model.

In this study the test animals were comprehensively tested for motor and selected non-motor features of PD followed by terminal histopathological analysis in the brain and the gut to confirm nigral degeneration of Tyrosine hydroxylase positive (TH+ve) neurons and Serine129 phosphorylated alpha synuclein aggregates (pSyn) pathology. We demonstrate that this 90 day P+L rat model of PD accurately simulates the temporal course of progressive Levodopa responsive motor deficits as seen in patients with PD^9^. These animals also exhibit non-motor symptomatology along with nigral degeneration and Psyn pathology in brain and gut.

This novel model therefore recapitulates the natural progression of the body first sub type of PD, accompanied by certain non-motor symptomatology thereby making it a robust model for preclinical evaluation of emerging therapeutic strategies.

## MATERIAL AND METHODS

All procedures were approved by the University of Toledo Institutional Animal Care and Utilization Committee (IACUC) and performed in accordance with the National Institute of Health guidelines.

### Materials

Unless indicated otherwise, all chemicals were obtained from Sigma-Aldrich (St. Louis, MO).

#### Animals and treatment

50 Male Sprague–Dawley rats weighing 200-300 g (7 to 9 weeks old) were housed in an AAALAC accredited Animal Care Facility maintained at 24 °C humidity on a 12:12 h light/dark cycle. Food and water were provided ad libitum. These animals were distributed into four distinct groups with each group receiving their respective treatment for ninety consecutive days as follows: Group 1 received Paraquat (0.078 mg/kg, p.o.) + Lectin (0.004%, p.o.) + Cholecystokinin (CCK) (3 µg/kg, i.p.) (P+L, n = 32), Group 2 received Paraquat (0.078 mg/kg p.o.) + CCK (3 µg/kg, i.p.) (P, n=6), Group 3 received Lectin (0.004%, p.o.) + CCK (3 µg/kg, i.p.) (L, n=6) and Group 4 received Sucrose solution (1%, p.o.) + CCK (3 µg/kg, i.p.) (Vehicle control/S, n=6). To promote absorption, gastric emptying was delayed by injection of cholecystokinin (3 μg/kg i.p.) 15 min prior to each gavage.

Group I was distributed into different cohorts and the animals were examined at different times in order to check the reproducibility and rigor of the experimental design and the results. A group of P+L+CCK rats (Group 1) received injections of L-dopa (4 mg/kg) and benserazide (15 mg/kg, i.p. diluted in ascorbate saline; n =5) twice a day for 2 days, after 90 days of P+L treatment.

### Tissue Collection

At the conclusion of the behavioral experiments, rats were euthanized under deep general anesthesia with ketamine and xylazine cocktail (0.2ml/100kg) followed by rapid sternal thoracotomy and transcardiac perfusion with 500 ml of room temperature followed by cold heparinized saline. The gastrointestinal tissue was collected at this stage for gut histology and then the animal was perfused with ice cold 4% paraformaldehyde (PFA) in PBS. Brains were isolated and postfixed in 4% PFA. Brains were transferred to a 15% sucrose solution in PBS followed by 30% sucrose solution in PBS for cryoprotection. Brains were sectioned in 60mm-thick coronal sections using a freezing microtome in 1:10 series and preserved in cryoprotectant until further processing.

### Gastrointestinal tissue collection

The gut tissue between oesophagus and the end of posterior colon was harvested after perfusion with ice-cold PBS. The proximal colon was separated and cleaned by flushing the interior with ice cold PBS thoroughly to remove all the fecal matter using a syringe. Once cleaned, the separation of longitudinal muscle and myenteric plexuses (LMMP) was carried out according to Huang et. al., 2021 ^10^. A 6mm glass rod was inserted into the lumen. A superficial incision was made along the serosal surface at the attachment line of the mesentery using a fine forceps. With the help of a cotton swab soaked in phosphate buffered saline, the strip of longitudinal muscle with myenteric plexus was gently peeled away. The stripped longitudinal muscle and the myenteric plexus (LMMP) were then placed on a supercooled polylysine coated glass slide, fixed using 4% paraformaldehyde for 20 minutes at 4°C followed by transfer into cold tris-buffered saline (TBS) buffer till staining.

### Behavioral testing

A well-established rodent behavioral battery of tests (RBBT) was used to identify the parkinsonian phenotype in treated rats, as described previously ^11^. Briefly, these consist of the

#### i Vibrissae-evoked forelimb placement test (VEFT)

Vibrissae test is a forced reflex test in which the tester restrains both hindlimbs and one forelimb of the rat, allowing one free forelimb. The rat is brought parallel to the surface of a table which allows stimulation of the ipsilateral vibrissae to evoke a reflex ipsilateral forelimb placement on the table surface. This test is repeated 10 × 3 at each testing session for each limb therefore a total of 30 vibrissae touches per limb. Average paw touches out of 10 trials were recorded. The test was performed at baseline (prior to treatment), 2, 4, 8 and 12 weeks of P+L+CCK treatment for vibrissae test, as well as 2 months after cessation of P+L+CCK treatment i.e. 21 week (before euthanasia)

#### ii Levodopa responsiveness

The motor impairment observed in the rats were evaluated for their levodopa responsiveness by administering L-dopa (4 mg/kg) and benserazide (15 mg/kg, i.p. diluted in ascorbate saline;) twice a day for 2 days after 90 days of P+L+CCK treatment ^12^. The rats were tested for the vibrissae score as mentioned in Section 1 of behavioral testing.

#### iii Cylinder test

The cylinder test was used to assess hindlimb and forelimb function, and protocol was adapted from those previously described ^11, 13–15^. The rat was placed into a transparent cylinder measuring (20 cm diameter, 25 cm high). Two mirrors were placed at a 90° angle from each other directly behind the cylinder and the video camera was placed in front of the camera to record all rears and paw touches. Rats were placed in the cylinder and video recorded for five minutes. The cylinder was cleaned with 70% ethanol after every test. Total right and left paw touches were then counted up to 20 total touches. A lateralization index was calculated, to analyze any right or left paw bias in the P+L treated rat. This was calculated by (left paw touches – right paw touches) / 20 paw touches, with a lateralization index of 0 meaning equal use of left and right paw, a positive number meaning a bias towards left paw and a negative number meaning a bias towards right paw touches^16–18^. The test was performed at baseline (prior to treatment), and 12 weeks of P+L+CCK treatment, as well as 2 months after cessation of P+L+CCK treatment i.e. 21 weeks (before euthanasia).

#### iv Y maze test

This test was performed to measure the short-term working memory in the test rats ^19^. The test rats were placed in a polycarbonate maze in the shape of a Y and allowed to roam freely for 5 minutes. Rats were tested with no previous exposure or habituation to the maze. Rat behavior was recorded and analyzed by a video tracking system by ANY-Maze. The total number of arm entries and the number of times alternates the arm were monitored and the percent of spontaneous alterations made by the rats was calculated. A spontaneous alternation was defined as an entry into three different arms on consecutive choices. The percentage of alternation was calculated as the ratio of actual to maximum number of alternations. The test was performed at baseline (prior to treatment), and 12 weeks of P+L+CCK treatment, as well as 2 months after cessation of P+L+CCK treatment i.e. 21 weeks (before euthanasia).

#### v Novel Object Recognition test (NORT)

The Novel object recognition test was performed to measure animals’ natural exploratory activity and the ability to recognize a novel object in the surrounding ^20, 21^. It consists of a 100 x 100 cm square arena in which the rats are allowed to explore the surroundings for 5 minutes. The behavior of rats was recorded and analyzed by a video tracking system by ANY-Maze. The NORT span over 3 days which includes familiarization, identical object and novel object days. On Day 1 i.e., familiarization, the rat is allowed to roam freely in the arena to get habituated to the NORT box. On day 2 i.e., identical object day, two identical objects are placed at opposite corners of the arena and the rats are allowed to explore. On day 3 i.e., Novel object day, one of the day 2 object is replaced by a novel object. The position of all the objects was kept constant throughout the experiment. After each trial the arena and object cues were cleaned with 70% ethanol to minimize olfactory cues. Working memory is evaluated by the discrimination index, which is defined as the time exploring the novel object divided by the total object exploration time. The test was performed at baseline (prior to treatment), and 12 weeks of P+L+CCK treatment, as well as 2 months after cessation of P+L+CCK treatment i.e. 21 weeks (before euthanasia).

#### vi Sleep analysis

To evaluate the effect on Rapid Eye Movement (REM) behavior the animal were video recorded over a period of 2 hours after the 90 day P+L treatment. The recordings were done in the lights on phase. Briefly, the animals were placed individually in a clean cage. They were provided with ad libitum food and water. Further, the animals were video recorded over a period of 2 hours for their sleep activity. The video recordings were scored for wakefulness, Non-REM (NREM) sleep, and REM sleep according to the behavior-based criteria for sleep-wake stage classification ^12^.

#### iv Immunohistochemistry

##### a. Tyrosine Hydroxylase (TH) staining

TH immunochemistry was done according to previously published protocol ^22, 23^. Briefly, 1:10 series of the whole brain free floating sections were washed extensively with PBS and permeated using PBS with 0.2% Triton-X100 (PBS-Tx). Post permeation sections were incubated in 0.3% hydrogen peroxide in PBS-Tx for 30 min at room temperature to inactivate endogenous peroxidase activity. Sections were blocked with 10% normal donkey serum in PBS-Tx for 1 hour at room temperature to block nonspecific binding sites and then incubated in primary antibody (rabbit anti-tyrosine hydroxylase, Pel Freez Biologicals #P40101-150, 1:250 in blocking solution) for 72 hours at 4°C on a shaker. Sections were washed in between incubation steps with 6 x 3min PBS-Tx. Sections were incubated in diluted secondary antibody (biotinylated donkey anti-rabbit IgG, Jackson ImmunoResearch #711-065-152, 1:200 in blocking solution) for 1 hour at room temperature. After washing sections were incubated with diluted streptavidin-horseradish peroxidase (ThermoFisher Scientific #21126, 1:500 in PBS) for 30 minutes at room temperature. Visualization of immunodetection was done with 3,3’-diaminobenzidine tetrahydrochloride (DAB) reaction for 1 min. Post development, the sections were washed and mounted on double gelatin-coated slides and dried for at least 12 hours. Sections were dehydrated in ascending grade ethanol concentrations (70%, 95%, 100%, 100%) for 3 min per ethanol and placed in histoclear until cover slipped with DPX mounting medium.

##### b. Phospho-S^129^-alpha synuclein staining

1:10 series of whole brain free floating sections were washed in tris-buffered saline (TBS) with 0.5% Triton-X100 (TBS-Tx) and incubated in 0.3% hydrogen peroxide to inactivate endogenous peroxidase activity. Non-specific binding sites were blocked with 10% normal goat serum in TBS-Tx for 1 hour and incubated in primary antibody solution (mouse anti-alpha synuclein phosphoS129, ab184674, 1:10000 in blocking solution) for 24 hours 4°C on a shaker. Sections were washed in between incubation steps with 4 x 5min TBS-Tx washes. Sections were then incubated with secondary antibody solution (Biotin-goat anti-mouse IgG Millipore AP124B, 1:500 in blocking solution) at 4°C on a shaker. After washes sections were incubated in ABC reagent (Vectastain Elite, PK-6100) for visualization of DAB reaction. After washing the developed sections were mounted on double gelatin-coated slides and dried for at least 12 hours. Sections were dehydrated in ascending grade ethanol concentrations (70%, 95%, 100%, 100% respectively) for 3min each and placed in histoclear until cover slipped with DPX mounting medium.

##### c. Proteinase K treatment

The methodology for Proteinase K treatment was adapted from the protocol by Patterson et. al. ^24^. Representative brain sections of a P+L-treated and naïve control animals containing the SNpc were washed in TBS. Sections were treated with 10µg/ml proteinase K (Invitrogen, 25,530,015) for 15 min at room temperature. Following proteinase K treatment, sections were washed in TBS and then TBS-Tx. Sections were incubated in 0.3% hydrogen peroxide. Sections were blocked with 10% Normal Goat Serum in TBS-Tx. Sections were incubated in primary antibody (anti-alpha synuclein, ab212184, 1:1000 in blocking solution). Sections were placed in secondary (goat anti-rabbit, ab205718, 1:500 in blocking solution). Sections were washed in between incubation steps with 4 x 5min washes with TBS-Tx. Visualization of immunodetection was done with DAB reaction. Sections were mounted on double gelatin-coated slides and dried for at least 12 hours. Sections were dehydrated in ascending ethanol concentrations (70%, 95%, 100%, 100%) for 3 min per ethanol and placed in histoclear until cover slipped with DPX.

##### d. Cresyl violet acetate staining

1:10 series of whole brain free floating sections stained for phospho-S^129^-a-synuclein were counterstained with cresyl violet acetate. Following immunostaining, sections were hydrated in descending ethanol concentrations (100%, 100%, 95%, 70%) followed by ddH^2^O prior to staining with 0.5% cresyl violet acetate (CV). Sections were then dehydrated with ascending ethanol concentrations (70%, 95%, 100%, 100%) and placed in histoclear until cover slipping with DPX. Representative 1:10 coronal sections of the entire brain stained with cresyl violet were examined to evaluate for any evidence of generalized toxicity.

##### e. Gut/Colon Phospho-S^129^-alpha synuclein staining

Following extraction of the myenteric plexus of the colon as stated abovethe tissue was washed in 0.5% Triton-100X in tris-buffered saline (TTBS) for 3 X 30 minutes, blocked in 0.3% Triton-100X in tris-buffered saline with 20% normal horse serum at room temperature on shaker for 2 hours, and incubated in primary antibody (chicken anti-beta-tubulin 3 (TUJ1; Aves laboratory; 1:1000) and mouse anti-alpha synuclein phospho-S129 antibody (Abcam, ab184674, 1:10000) in 0.5% Triton-100X in tris-buffered saline and 20% horse serum on shaker at 4°C for 72hrs. Tissue was washed in 0.5% Triton-100X in tris-buffered saline and 20% horse serum 3 X 1 hour, washed in 0.5% Triton-100X in tris-buffered saline 3 X 10 minutes, and incubated in secondary antibody (donkey anti-mouse-594 1:1000 and donkey anti-chicken-488 1:1000) in 0.5% Triton-100X in tris-buffered saline on shaker at 4°C for 2 days. Tissue was washed extensively with 0.5% Triton-100X in tris-buffered saline 4 X 10 minutes, mounted using Vectashield hardset (Vector laboratories) with DAPI and visualized for fluorescence.

#### v Stereology

A 1:5 series of brain sections which contain the whole SNpc were stained for TH via immunohistochemistry as described above. TH+ neurons were quantified using Stereo Investigator software suite from MicroBrightfield (MBF) bioscience with a 100X magnification using an Olympus BX53 microscope (Olympus, Tokyo, Japan) fitted with a digital CCD camera (Hamamatsu, Hamamatsu City, Japan) and a motorized stage (Prior Scientific, Rockland, MA, USA). The total number of TH+ neurons in the SNpc was estimated using design based optical fractionator method of stereology. The coefficient of error (CE) was calculated according to Gundersen et. al., ^25^and values were accepted if CE £ 0.2. TH-stained sections were counterstained with CV to differentiate between TH downregulation and neuronal loss.

#### vi Phospho-S129-alpha synuclein pSyn quantitation

1:10 series of 60µ sections were stained for pSyn as described above. Slides were scanned with a slideview VS200 slide scanner and viewed with OlyVia imaging software. ROIs were drawn around the SNpc at 10X magnification and markers placed on observable pSyn aggregates at 40X magnification.

### Statistical analysis

Data are reported as mean ± SEM and in all instances, significance was set at p < 0.05. Data was evaluated using or Two-way ANOVA repeated measures mixed effect model followed by Bonferroni’s multiple comparison test for vibrissae test and Two-way ANOVA followed by Bonferroni’s multiple comparison test for levodopa responsiveness and cylinder using GraphPad® software (Graph Pad Prism). Stereology counts of TH+ neurons was evaluated for significance using independent t tests in R. Graphs were created with GraphPad Prism software.

## RESULTS

### A. Behavioral tests

#### i. Chronic P+L administration leads to the induction and gradual progression of motor impairment in study animals

The VEFT revealed a slow gradual onset as well as progression of the motor impairment in the animals which received P+L (Figure.1) as compared to the control animals. Figure. 1 shows the timeline for the first cohort (n=8) of the Group I. The onset of motor deficit was observed after 4 weeks (4 weeks-Left: 8.77±0.23, Right: 8.88± 0.11 vs Baseline-Left: 10± 0.0, Right: 9.98±0.01) of P+L treatment with the development of hemiparkinsonian (unilateral) symptoms in about 25% of animals while the rest 75 % still showed no effect. The symptoms in these animals proceeded to deteriorate progressively with the animals developing bilateral symptoms around 12 weeks of P+L treatment (8 weeks-Left: 7.12± 0.13, Right: 7.80± 0.10 vs Baseline-Left: 10, Right: 9.98±0.01; 12 weeks: Left: 2.37± 0.15, Right: 6.07± 0.14 vs Baseline-Left: 10± 0.0, Right: 9.98±0.01). The progression in the severity of the symptoms was observed even after cessation of the treatment and interestingly, at the end of 21 weeks of the study duration, three distinct subsets of animals emerged with regards to the severity of symptoms. At the end of 21 weeks the P+L cohort consisted of 50% bilateral, 25 % unilateral/hemiparkinsonian and 25% animals with no symptoms. This was evident by the significant gradual consistent decline in the vibrissae score for left and right forelimb (21 weeks-Left: 2.24± 0.04, Right: 4.50± 0.04) in the P+L treated animals as compared to their baseline score as well as with that of control (21 week-Left: P (10± 0.0), L (10± 0.0), S (10± 0.0), Right-P (10± 0.0), L (10± 0.0), S (10± 0.0)) at the specific time points. Also, no P+L treated animal exhibited spontaneous recovery of motor impairments over the duration of the study. The controls did not significantly differ in their vibrissae score over the entire duration of the experiment (Figure. 2).

**Fig. 1.:**
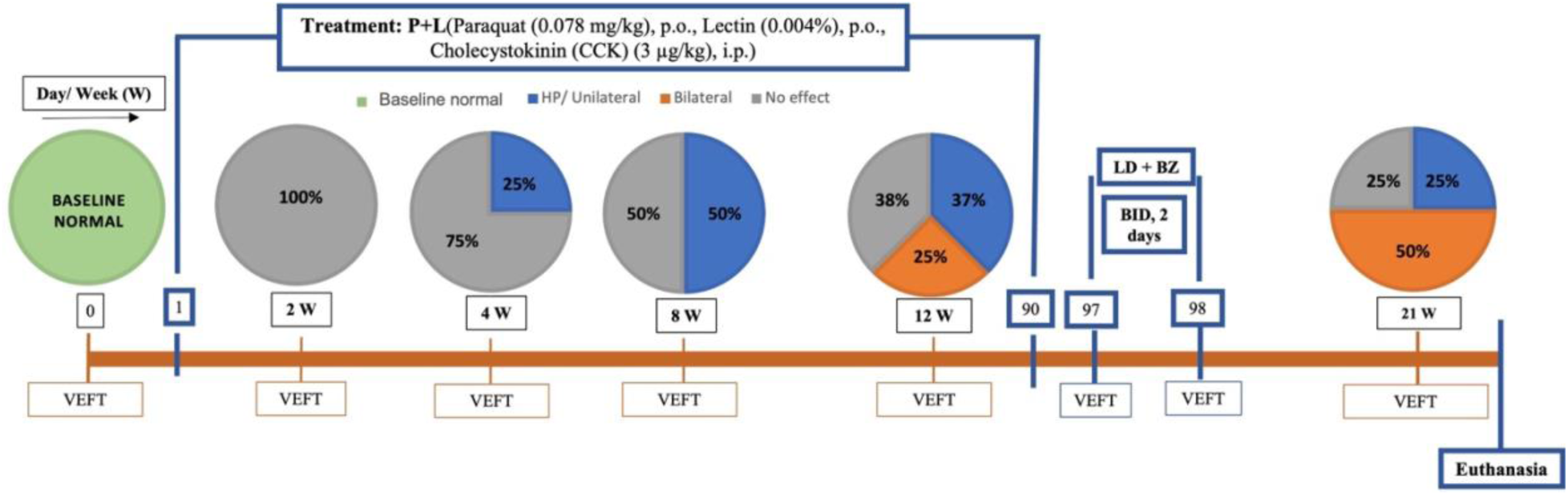
Experimental timeline and schematic representation of the evolution of parkinsonian features in the P+L treated animals based on Vibrissae test over the duration of the experiment (n=8). VEFT: Vibrissae tests, W: Weeks, %: Percentage of animals affected, LD+BZ: Levodopa (4 mg/kg) + Benserazide (15 mg/kg, i.p.) twice a day (BID) for two days.

**Fig. 2.**
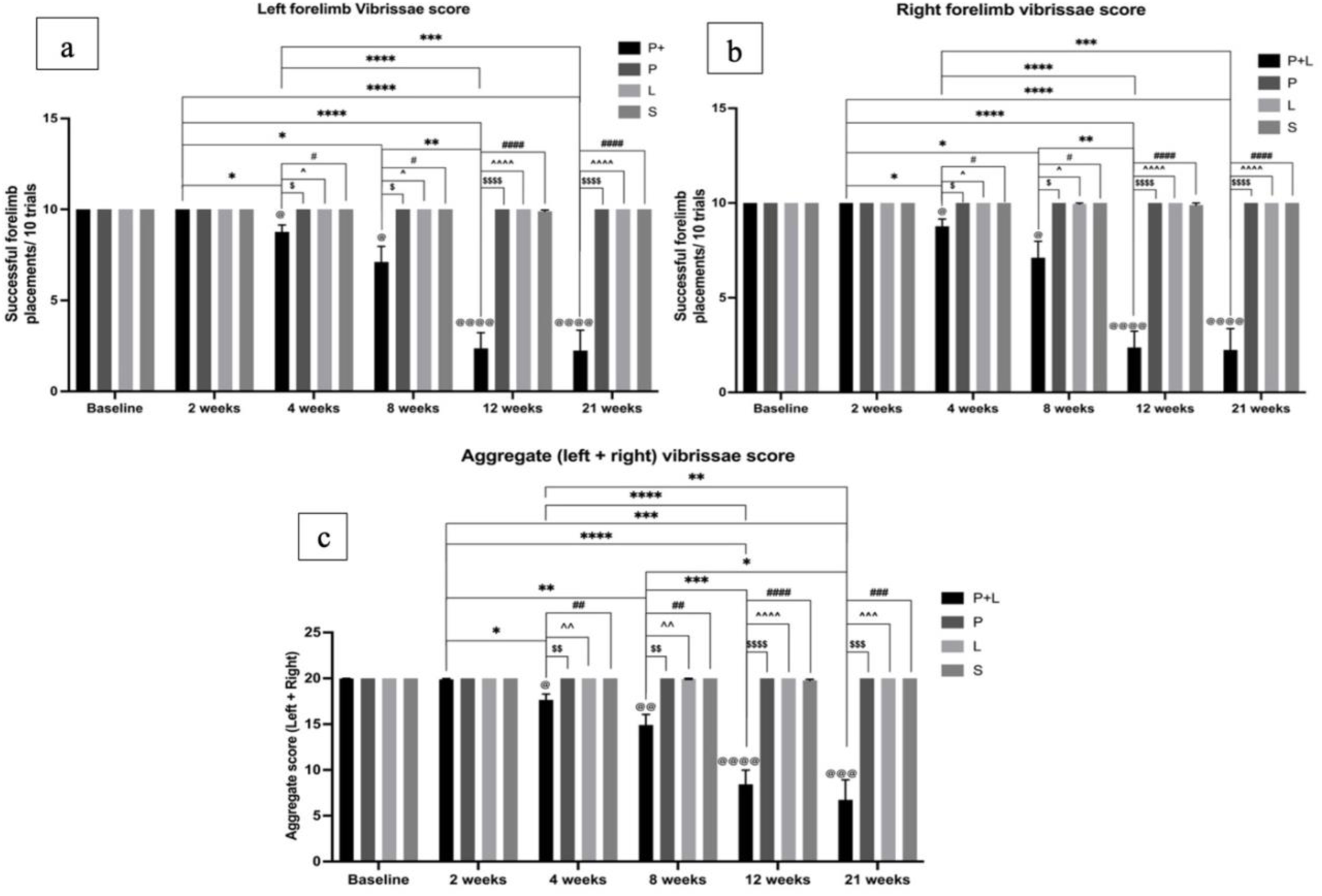
Vibrissae score over the entire duration of the experiment. a. Vibrissae score of left forelimbs of test animals representing the motor deficits in left forearms over the duration of the experiment. b. Vibrissae score of right forelimbs of test animals representing the motor deficits in right forearms over the duration of the experiment. c. Aggregate vibrissae (left + right) score representing the composite overall motor deficit in both arms over the duration of the experiment. Data expressed as mean ±SEM. P+L n=32; P, L and S n=6. The data was analyzed by Two-way ANOVA repeated measures mixed effect model followed by Bonferroni’s multiple comparison test. *, $, ^, # p<0.05, **, $$, ^^, ## p< 0.01, ***, $$$, ^^^, ### p<0.001 and @@@@, ****, $$$$, ^^^^, #### p<0.0001 vs baseline P+L, P+L P, L and S respectively.

#### ii. Levodopa responsiveness in vibrissae test

An expected improvement in the motor performance in the P+L animals consisting of three hemiparkinsonian (Unilateral deficit) and two bilaterally affected animals, was observed after L-Dopa treatment (Levodopa (4 mg/kg) + Benserazide (15 mg/kg, i.p.) twice a day (BID) for two days). The amelioration in the parkinsonian symptoms was apparent by the significant increase (p<0.01) in the cumulative vibrissae score of the P+L animals after treated with 4 doses of Levodopa.

**Fig. 3.**
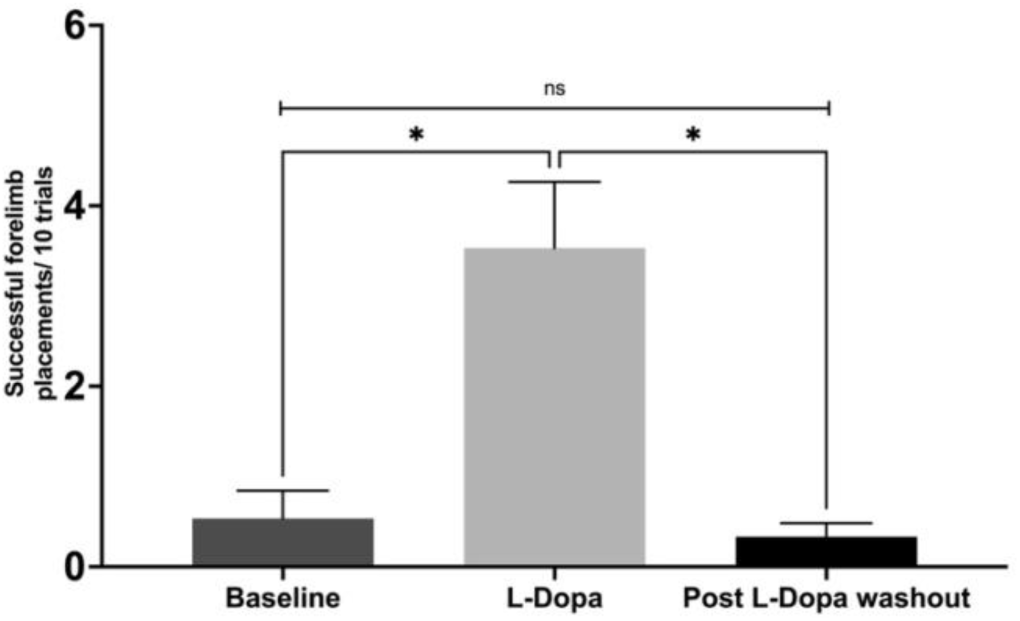
Levodopa responsiveness (Levodopa (4 mg/kg) + Benserazide (15 mg/kg, i.p.) twice a day (BID) for two days) in vibrissae test. Data expressed as mean ±SEM, n=5. **p<0.01 using One-way ANOVA followed by Bonferroni’s multiple comparison test.

#### iii. Cylinder test

The cylinder test was used to investigate if the motor defects were manifested more in a particular arm than the other as an indicator of asymmetry. To determine this, we assessed the lateralization index of the P+L treated rats in the cylinder test at the baseline, 12 weeks and 21 weeks of the study. The lateralization index ranged between +1 to -1 wherein the positive number indicates more use of left paw, negative number indicates more use of right paw and a 0 would indicate equal use of both paws ^16–18^. As seen in Figure.4a, at 12 weeks the lateralization index leans more negatively towards right paw use as compared to the baseline and 21 weeks. This indicates that motor deficits in majority of animals in this cohort are more apparent in left paw use which is displayed as higher motor asymmetry at 12 weeks. This finding is further substantiated with a relative greater percent of decrease in the number of touches by left forepaw noted at 12 weeks with respect to baseline readings as compared to the right forepaw (Figure.4b). A lower lateralization at 21 weeks is due to the higher number of bilaterally affected animals at this stage in this model.

**Fig. 4.**
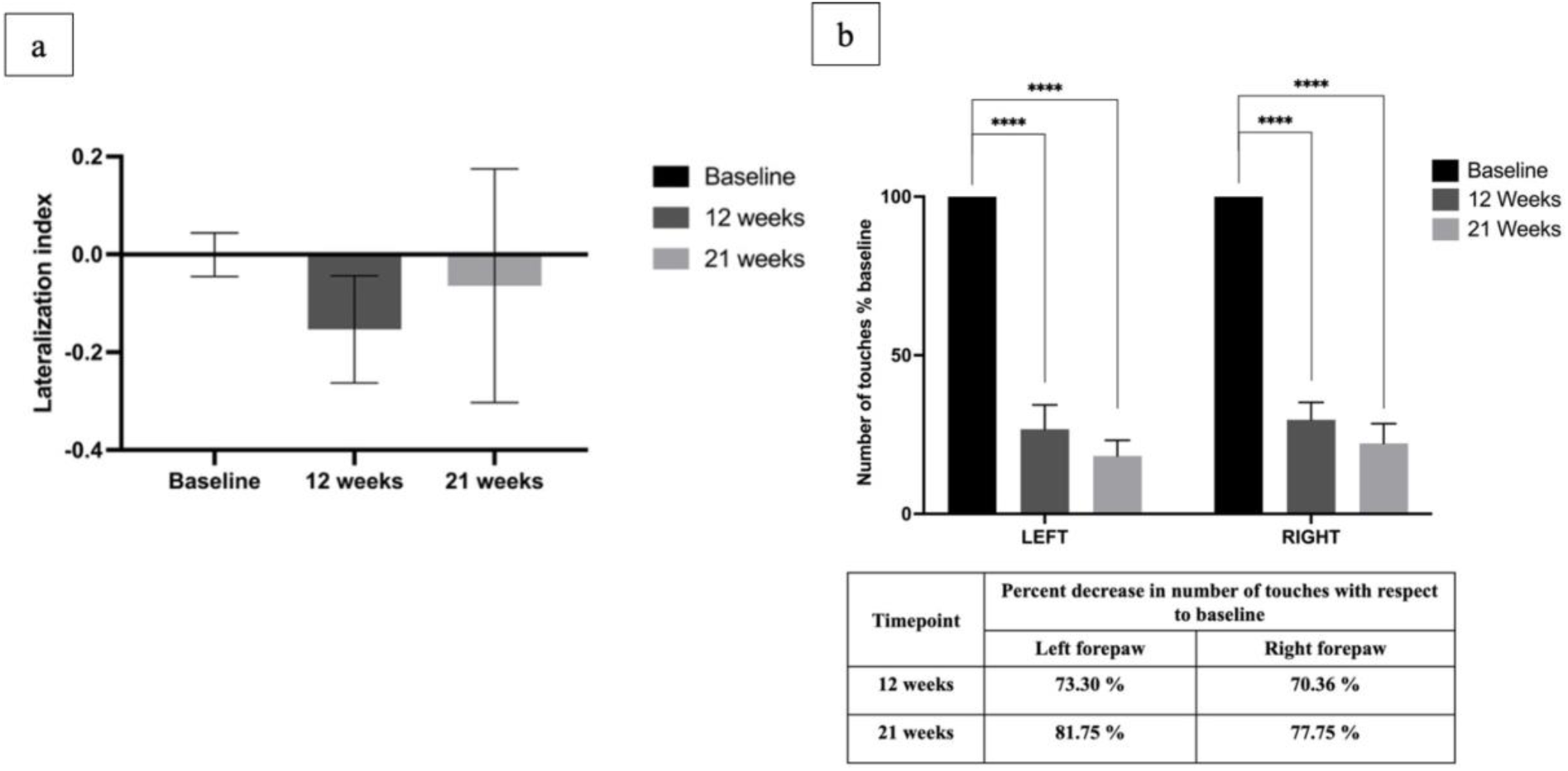
Cylinder test for assessing the limb asymmetry. a. Lateralization index over the duration of the study. Data expressed as mean ±SEM. The data was analyzed by One-way ANOVA repeated measures mixed effect model followed by Bonferroni’s multiple comparison test. b. Percent decrease in the number of left and right touches with respect to baseline readings in P+L treated animals. Data expressed as mean ±SEM. ***p<0.001, ****p<0.0001 using Two ANOVA test followed by Bonferroni’s multiple comparison test.

##### iv. Y maze test

The animals were subjected to Y maze test in order to analyze the effect of P+L treatment on their short-term working memory. The cognitive deficit in the animals was evaluated by measuring the spontaneous alternations made by the animals with in the three arms of the Y maze. As evident from Figure. 5 the P+L treated animals exhibited a significant decrease in the percent spontaneous alteration as observed at 12 weeks of the P+L treatment (p<0.001 vs baseline P+L, 12-week L, S and p<0.05 vs 12-week P). The decline was evident even 2 months after the cessation of the treatment i.e. 21 weeks timepoint (p<0.0001 vs baseline P+L, p<0.05 vs 21 weeks P, p<0.01 vs 21 weeks L and S). The control groups including P, L and S did not show any significant difference in the percent spontaneous alterations during the entire duration of the experiment.

**Fig. 5.**
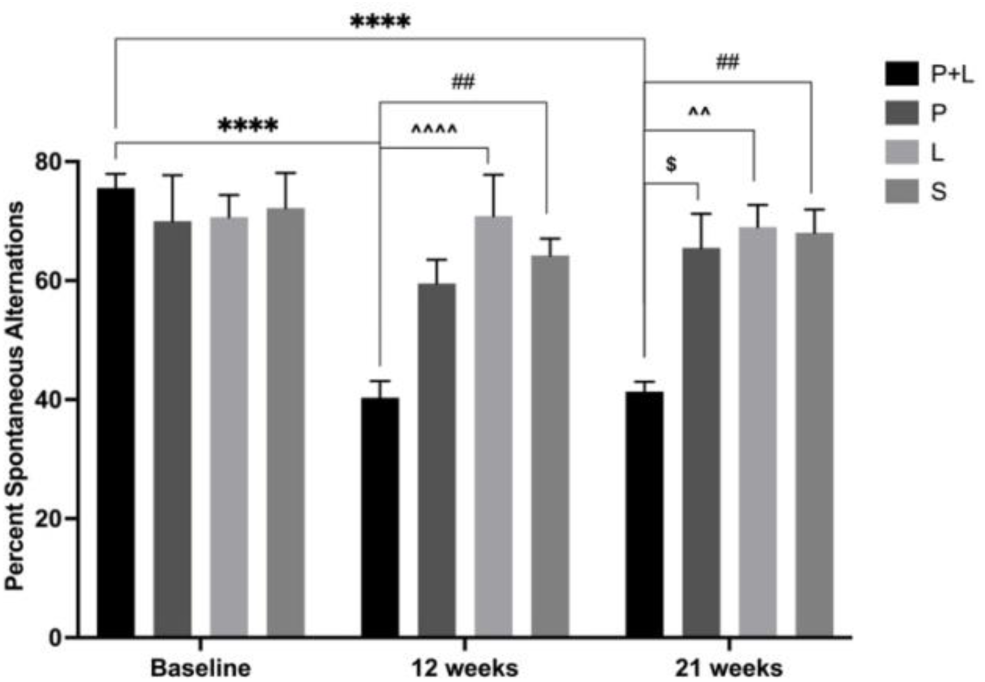
Percent spontaneous alteration in Y maze test over the entire duration of the experiment. Data expressed as mean ±SEM. The data was analyzed by Two-way ANOVA followed by Bonferroni’s multiple comparison test. *, $, ^, # p<0.05, **, $$, ^^, ## p< 0.01, ***, $$$, ^^^, ### p<0.001 and ****, $$$$, ^^^^, #### p<0.0001 vs baseline P+L, P, L and S respectively.

#### v. Novel object recognition test (NORT)

The animals were subjected to assess the acquisition and retention of long-term memory. The activity of the animal in this test depends on their natural tendency to explore the novel object more than the familiar ones and is a measure of their recognition memory ^21, 26^. The long-term memory of the test animals was measured by their discrimination index based upon their ability to differentiate between and the amount of time spent near the novel and familiar object. As seen in Figure. 6., a significant decline in the discrimination index was observed in the P+L animals at the 12 weeks after the initiation of the treatment (p<0.001 vs baseline P+L, p< 0.01 vs 12-week P, L and S). A progressive decline in the discrimination index was observed at 5 months even after the termination of the treatment at 90 days (p< 0.0001 vs baseline P+L, 21 weeks P, L and S. The animals in the all the control groups including P, L and S retained a consistent discrimination index without aby significant changes during the entire duration of the experiment.

**Fig. 6.**
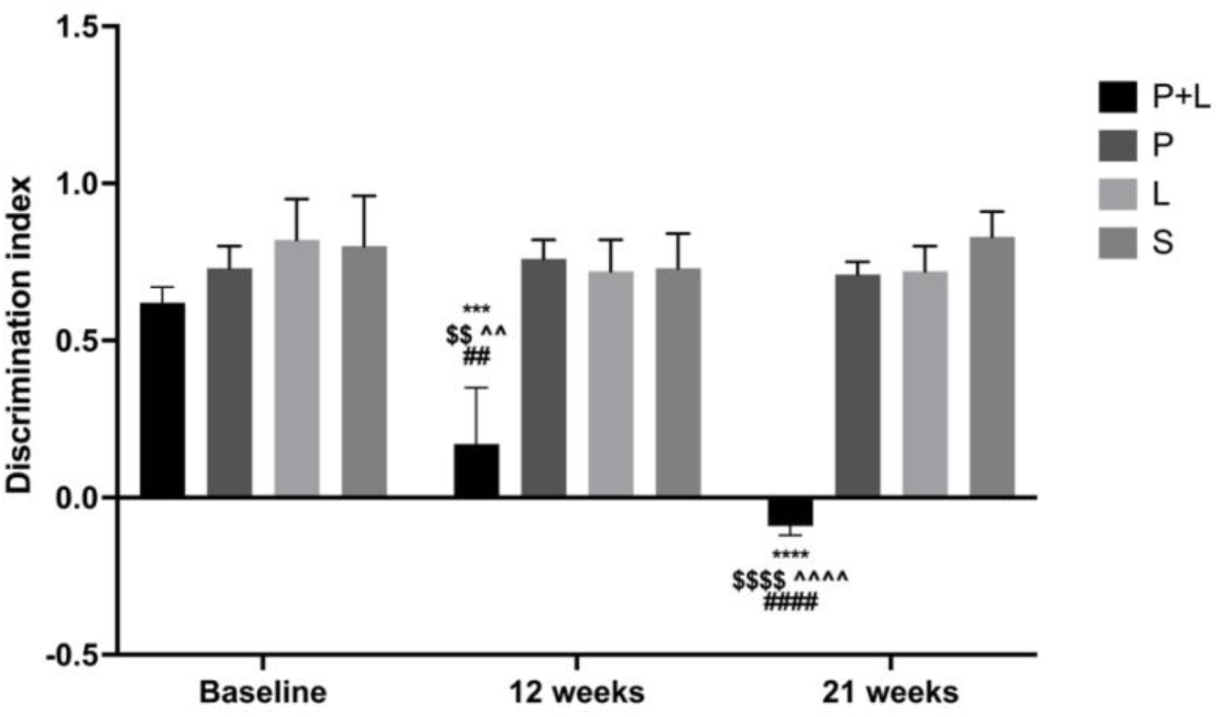
Discrimination index of the test animals in Novel object recognition test over the entire duration of the experiment. Data expressed as mean ±SEM. The data was analyzed by Two-way ANOVA followed by Bonferroni’s multiple comparison test. *, $, ^, # p<0.05, **, $$, ^^, ## p< 0.01, ***, $$$, ^^^, ### p<0.001 and ****, $$$$, ^^^^, #### p<0.0001 vs baseline P+L, P, L and S respectively.

#### vi. Sleep analysis

The sleep study recording exhibited altered REM-sleep behavior in P+L treated animals. This was evident by a significant difference in the time spent by the animals in the REM sleep stage when compared with naïve control animals (Figure 7. *p< 0.05 vs control, **p<0.01 vs control). This REM-sleep behavior disorder is observed in all three groups of PD symptoms severity despite the variations in the effect. No significant difference was observed in the NREM and wake sleep time in animals with different PD severity groups as well as when compared with the naïve control group.

**Fig. 7.**
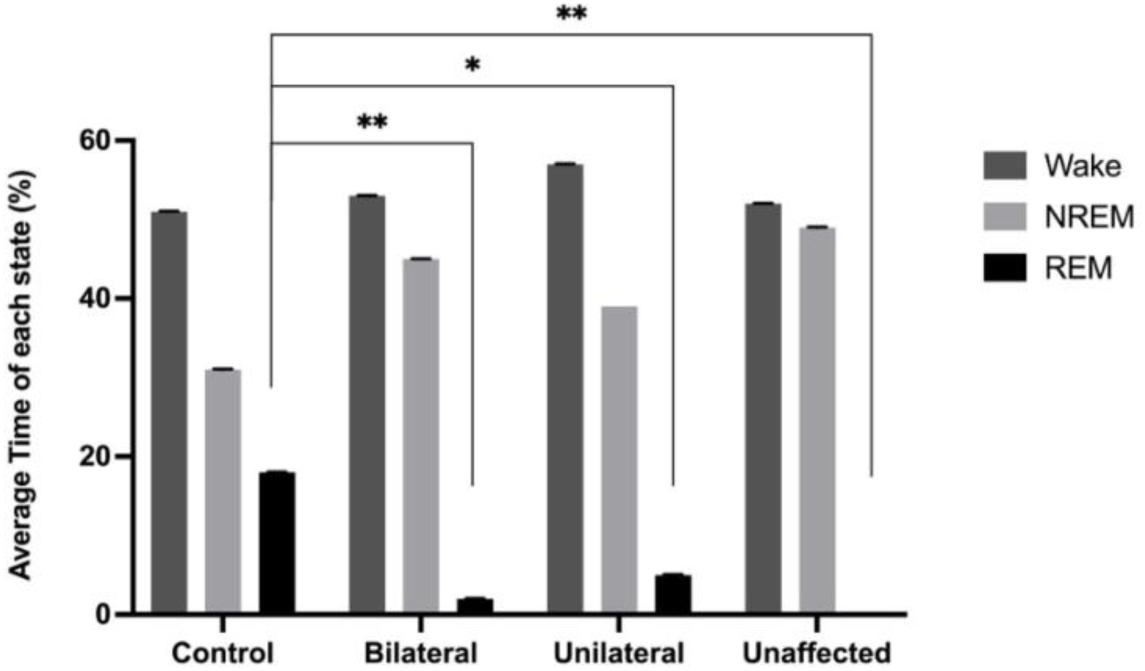
The effect of P+L administration on the average time spent by animals with different symptom severity in different sleep stages. N=8 Data expressed as mean ±SEM. *p<0.05 vs. baseline and **p<0.01 using One way ANOVA followed by Bonferroni’s multiple comparison test.

#### vi. Immunohistochemical analysis

##### a. Tyrosine Hydroxylase (TH) positive neurons in SNpc

The SNpc of the animals in different subsets of P+L and control groups were qualitatively compared for the neuronal loss in the TH positive neurons. As seen in Figure 8a, a discernible loss of TH positive neurons is seen in both SNpc of bilaterally affected animals while unilaterally affected animal exhibits an evidently greater loss in the contralateral i.e. right SNpc. The unaffected animals (no effect) show qualitatively equal amount of TH positive neurons in both SNpc. The stereological estimates (Figure 8b) of TH positive neuronal SNpc count in the rats with bilateral effects confirms the qualitatively findings and was found to be significantly different (p<0.001) as compared to the normal control rats.

**Fig. 8.**
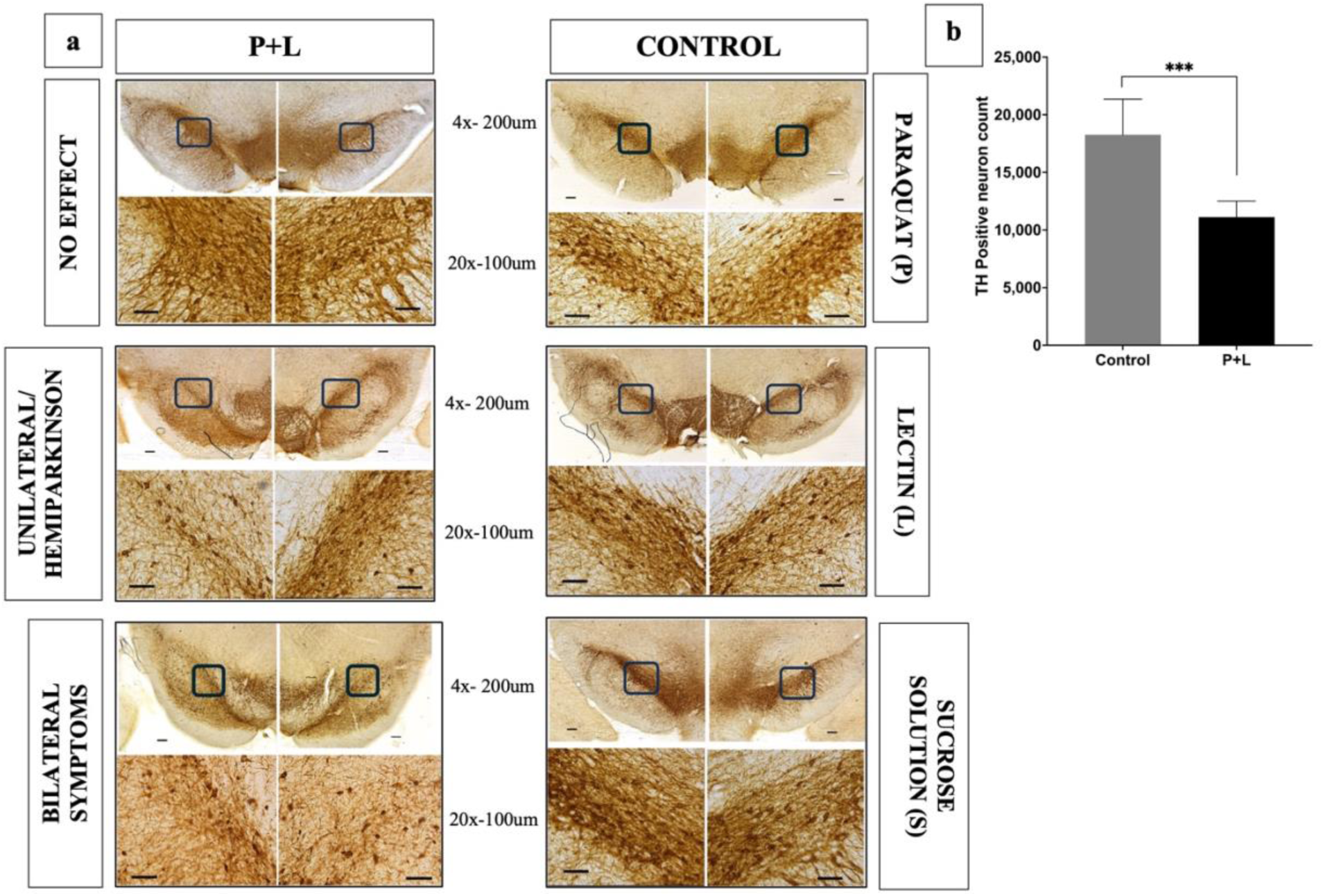
Effect of different treatments on TH positive dopaminergic neurons in SNpc. a. Representative images of TH-positive neurons in SNpc in different study groups. b. Stereological quantification of bilateral Tyrosine hydroxylase positive SNpc neuronal count in bilateral rats (n=4) compared to healthy untreated control rats (n=7). Data expressed as mean ±SEM. ***p<0.001 vs. normal control using independent t-test.

##### b. Phospho-S129-alpha synuclein (pSyn) and Proteinase K resistant alpha Synuclein

The SNpc region was qualitatively and quantitatively assessed for the presence of Phospho-S129-alpha synuclein. As seen in Figure 9a, both right and left SNpc of animals with bilateral motor deficit, have stained positive for Phospho-S129-alpha synuclein (pSyn). The presence of α-synuclein was apparent in the SNpc of the animals with unilateral as well. The a-synuclein aggregates were characteristic of Lewy body pathology as evidenced by their property of being proteinase K resistant as seen in Figure 9b. The quantitative assessment of pSyn revealed a significant increase in the pSyn count in the bilateral animals (*p<0.05) when compared with. No effect animals. The unilateral animals did show a significant increase in the pSyn count when compared with the no effect subgroup (Figure 9c).

**Fig. 9.**
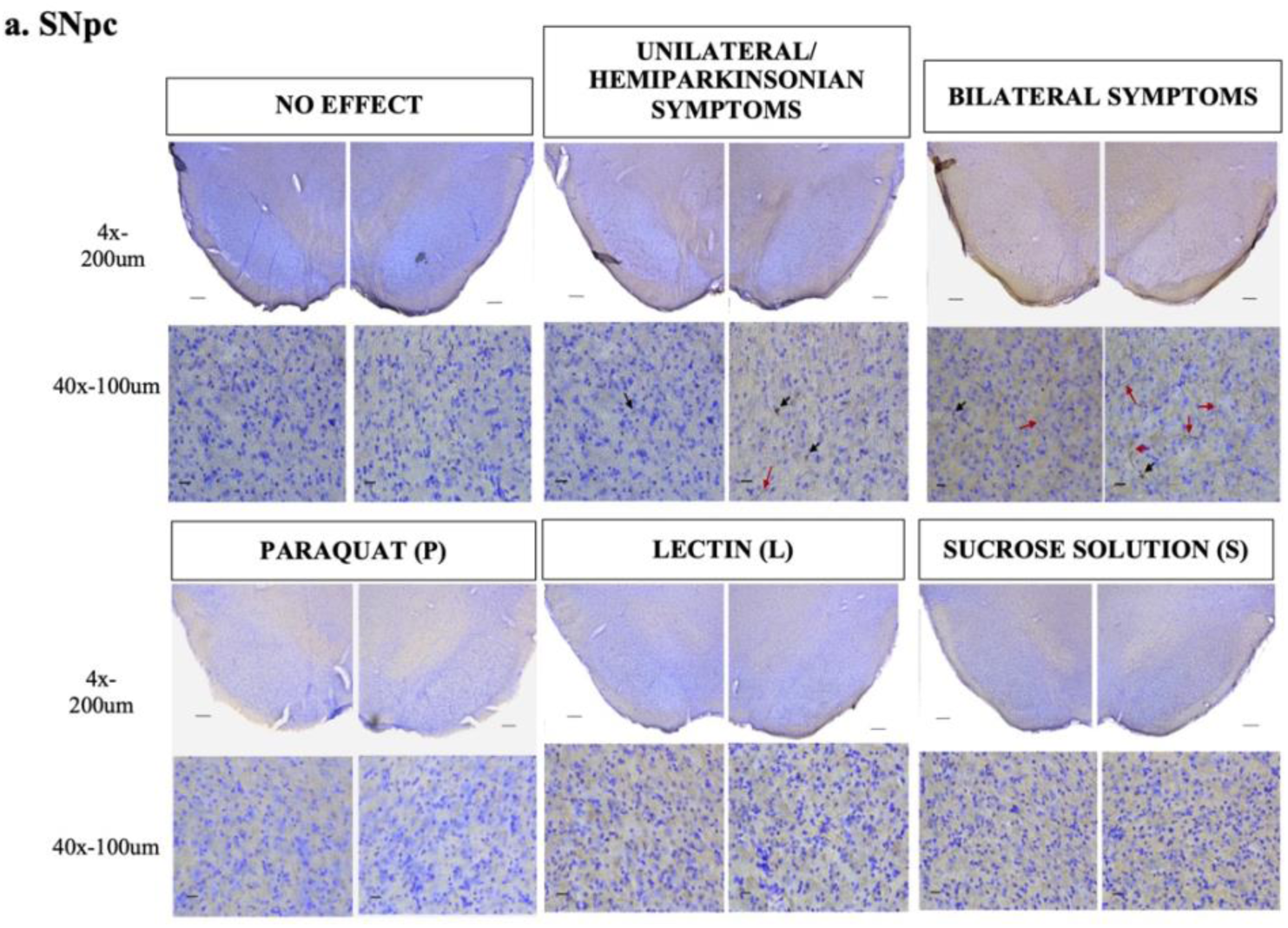
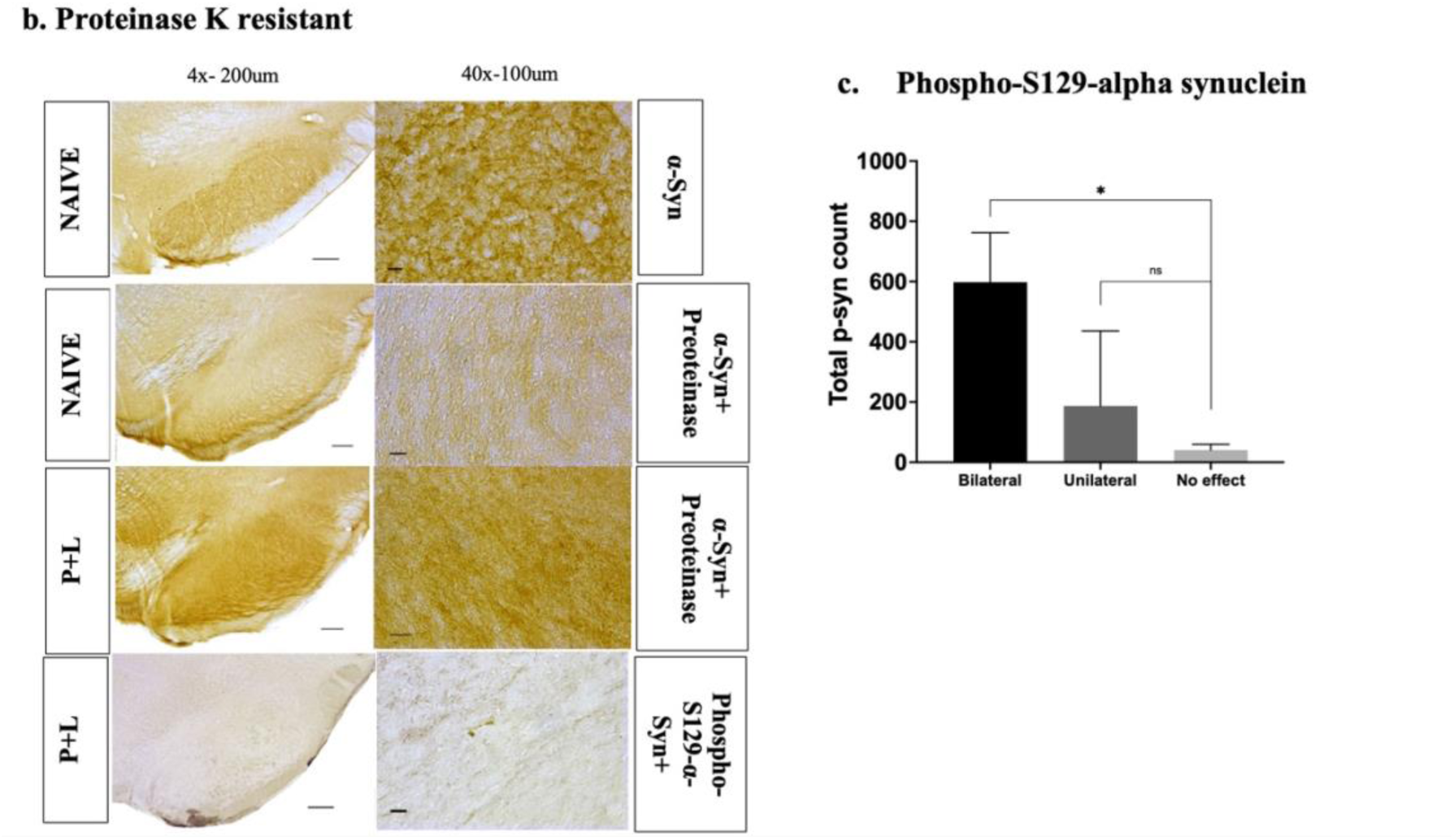
Phospho-S129-alpha synuclein immunostaining of different treatment groups. a. Representative images of Phospho-S129-α-synuclein immunostaining of SNpc of different study groups (Black arrow: α-synuclein aggregates and red arrows: α-synuclein fibrils) b. Proteinase K (PRK) resistance in naïve control vs P+L group. c. Stereology for total p-syn count in bilateral rats compared to no effect rats. n=2. Data expressed as mean ±SEM. *p<0.05 vs. no effect using independent t-test.

##### c. Gut/Colon Phospho-S129-alpha synuclein staining

The myenteric neurons in the rat colon was qualitatively evaluated for the presence of Phosphorylated S129-alpha synuclein (pSyn). The P+L treated animals at the end of the experiment showed the presence of pSyn while the 129Ser α-synuclein-immunoreactivity was not observed in naïve control animals. Therefore, the histopathological presence of pSyn aggregation was noted in the P+L treated animals.

**Fig. 10.**
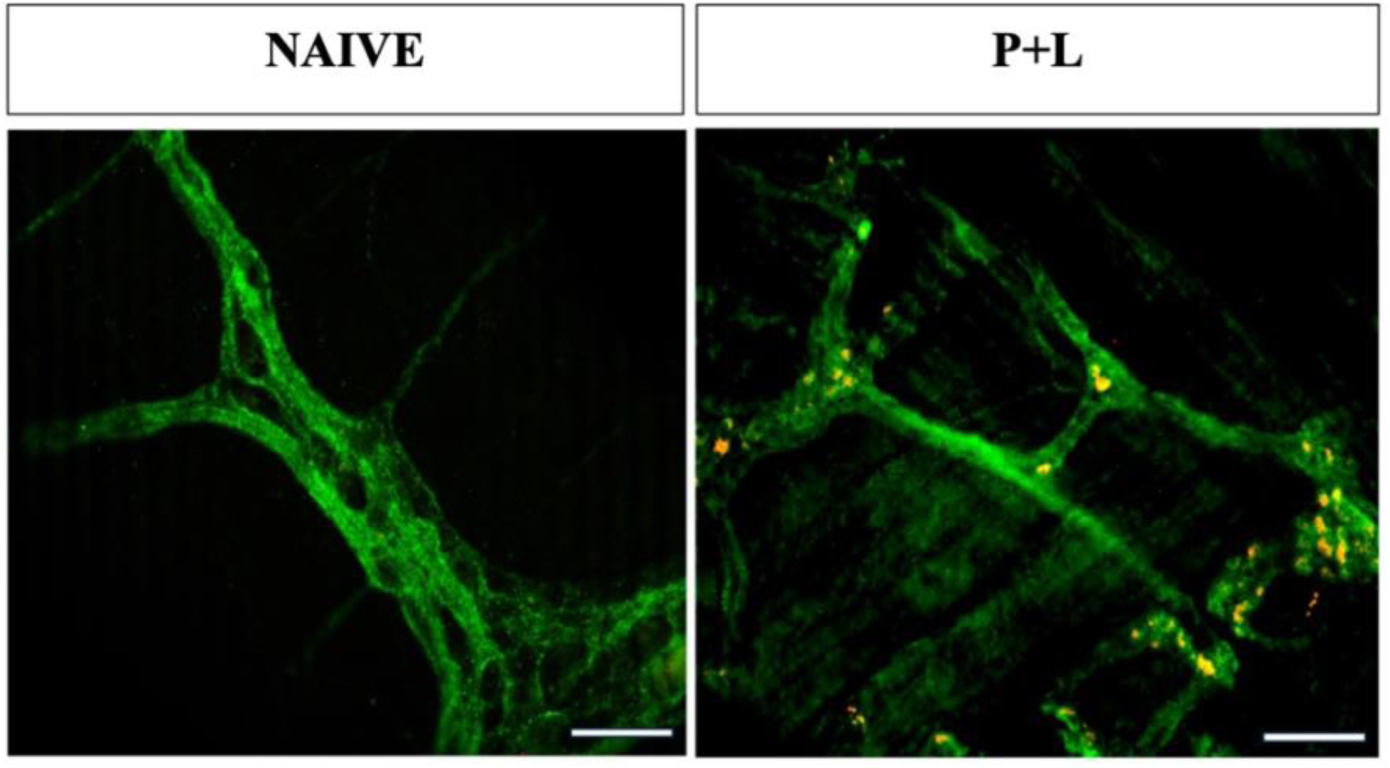
Myenteric plexus of colon stained for TUJ1 Beta Tubulin (green), phospho-S129-alpha-synuclein (red) in P+L and Naïve control rats.

## Discussion

The present study successfully demonstrates that the low daily dose of Paraquat and Lectin over a period of 90 days is able to replicate the natural course of idiopathic body first sub-type of PD. This dosage regimen leads to induction of characteristic onset of L-dopa responsive hemiparkinsonism in test animals that progressed to stable bilaterality over time. These progressive motors symptoms were accompanied by notable cognitive deficits as well. In addition, the terminal pathology exhibited the hallmark nigrostriatal dopaminergic degeneration accompanied by phosphorylated Serine129 Synuclein positive inclusions and evidence of gut pathology. Finally, we also demonstrate presence of sleep abnormalities that are being implicated in this sub type of PD therefore fulfilling a critical gap in PD research.

Parkinson’s disease (PD) is a result of a complex multifactorial pathophysiological dysfunction. It is now known to be a heterogenous disease with variability in pathological progression and clinical symptom progression ^27, 28^. Based upon this heterogeneity, researchers have in part differentiated PD into two subtypes which includes the “body-first” and “brain-first” PD. This classification is based on the differences in the site of initiation of disease, the progression pattern of alpha synuclein pathology, the early prodromal symptoms and the rate of progression of symptoms^1, 4, 5, 29^. As the name suggests the body first subtype initiates in the GIT and progresses retrogradely via vagus to SNpc exhibiting gut to brain progression. While in the brain first PD the α-syn pathology initiates in the brain and descends to the GIT. Researchers are still delineating the extent of the differences but the known distinction in the pathophysiology of PD advocates the need for tailored subtype specific therapeutic approaches. The preclinical experimental animal models which specifically recapitulates these distinct pathophysiological subtypes would prove to be an excellent platform to validate and translate these differences as well as to evaluate targeted novel therapies.

Currently available PD animal models including the neurotoxin and genetic models have their own advantages and disadvantages, but nearly all of them fail to completely mimic the pathophysiology of PD and not yet have been classified from an PD subtype perspective. Considering the existing data, historically idiopathic PD research was always aimed at nigrostriatal dopaminergic degeneration, the extranigral pathologies were overlooked while designing of the model. The available neurotoxin models either need intracranial administration of the toxin such as with 6-OHDA or the systemically administered toxins such as MPTP, rotenone, paraquat etc. do not replicate the route of administration. These models use a high dose of toxin to induce severe dopaminergic degeneration and parkinsonism which do not follow the temporal pattern and natural course of the disease. Since most of them mimic the nigral dopaminergic degeneration these can be classified as the brain first animal models ^29^. Though, past few years have seen a considerable improvement in the design of preclinical models, none of them till this date successfully simulates the full repertoire of symptomatology and pathogenesis of PD.

To overcome this, our lab previously has described a subthreshold P+L 7-day exposure animal model which replicates the route of environmental toxin entry as well as mimics body first subtype of PD as confirmed by bilateral sub-diaphragmatic vagotomy ^7, 30^. This 7-day model has a temporal course with disease onset within 2 weeks and complete evolution of motor dysfunction within 4 weeks. Recently we have also provided evidence of late cognitive deficits in this model ^30^. One of the major limitations of this model has been its shortened temporal course which does not allow the gradual progression of motor and non-motor symptoms to be studied in a slow, progressive manner, as observed in PD patients. Therefore, in the present study we have made an attempt at improving upon this shortcoming with designing and characterizing a progressively degenerative chronic rat P+L model. This chronic model constitutes repeated exposure of low daily doses of P+L while maintaining the total cumulative dose of 90 day same as in the 7-day model.

Our data from the 90-day daily low dose P+L oral exposure model in the rat shows that it faithfully replicates the natural progression of PD. The chronic exposure paradigm used in our study resulted in a gradual progression of the motor symptoms starting with a significant asymmetry. The symptoms initiated with the unilateral motor dysfunction at 4 weeks and eventually worsened into bilateral symptoms in about 50% of animals towards the end of the experiment. This is in line with the clinical observations where in human PD is considered to be an asymmetrical condition and in early stages it emerges majorly as an unilateral condition (Stage I hemiparkinsonism) ^9, 31–33^ and then advancing to stage II (bilateral disease) as originally defined by Drs. Hoehn and Yahr. It is also observed that although the symptoms eventually progress bilaterally, the initial side of onset might still exhibit more severe symptoms throughout the disease progression^33^. As apparent in our VEFT and laterization index results, the severity of the symptom in the left forelimb of the affected animals was greater than that of the right forelimb throughout the study duration. It is also important to note that the symptoms continued to evolve even after cessation of the exposure to Paraquat, suggesting a long-term effect of Paraquat exposure. Interestingly, in VEFT some of the animals were found to be unaffected over the entire period of the study and these constituted to 25% of total animals. This can be attributed to the variability in the susceptibility in the animals towards environmental toxin exposure similar to that observed in humans ^34–36^. Therefore, at the end of the experiment we achieved animal subsets with different motor symptom severity. Our model replicates a natural course of the disease progression which is stable and without any spontaneous recovery as observed throughout the experimental timeline of the study. To our knowledge this is the first model of PD that replicates this natural progression.

Further, we characterized the chronic exposure paradigm for its ability to induce non-motor symptoms specifically cognitive deficits and sleep abnormalities in these animals. As apparent from the results a significant progressive decline in the working short-term as well as long term memory was evident in the P+L treated animals indicating a pattern of progression in terms of cognitive dysfunction. The model was also able to produce sleep abnormalities regardless of the motor severity in the animals. Our model therefore was able to recapitulate selected non-motor symptomatology of PD.

The loss of SNpc neurons and their striatal projections in these animals compared to matched controls further provide proof of specificity of the model. The low dose of P+L administered orally daily for 90 days makes the daily dose very similar to what is likely to be representative of the exposure of P+L to humans prior to developing PD. Given that the average lifespan of laboratory rats in controlled environment is about 2 years, 90 days of exposure can be approximated to about 10 years in the human (1/8^th^ of 80 years). This approximation is relevant because in most situations P+L exposure in the human is not daily and is likely spread out in time, but likely cumulatively adding up to approximately 10 years.

One of the characteristic neuropathological features of PD is the cytoplasmic accumulation of Phospho-S129-α-synuclein as Lewy bodies and Lewy neurites in the surviving dopaminergic and non-dopaminergic and enteric neurons. In the present study, immunohistochemical analysis showed the presence of proteinase K resistant Phospho-S129-α-synuclein in both right and left SNpc of unilateral and bilateral animals as well as myenteric neurons in the colon stained positive for pSyn, therefore, providing evidence of α-synucleinopathy in the gut brain axis. We have shown in our 7-day model that bilateral truncal vagotomy prior to P+L exposure protects the rat from parkinsonism providing proof that the oral exposure to P+L triggers parkinsonism that is mediated by retrograde progression of alpha synuclein aggregation to the brain via the vagus nerves in the P+L rat model therefore, signifying the body first mechanism of progression.

Based on the imaging data of the prodromal and diagnosed PD patients, attempts have been made to stratify the PD subtypes based on the non-motor symptoms since the variability in them may correlate with the initiation and development of the α-synuclein pathology ^1–3, 29^. One of the important distinctions between the body and brain first subtype of PD has been noted as the presence of non-motor symptoms, particularly RBD in the early prodromal stages ^1, 4, 37^. As observed, our present chronic model shows a longer prodromal phase- and spaced-out early PD with 50% of unilateral animals achieved at only 8 weeks during the P+L exposure. Therefore, this novel model opens the window of time during which the phenotypic evolution of this sub type is faithfully captured allowing evaluation of targeted disease modifying therapies.

Altogether, the chronic exposure of subthreshold doses of Paraquat in combination with lectin and CCK is able to recapitulate the cardinal characteristic features of Parkinson’s disease including the variability and progression of motor impairment, responsiveness of the observed motor impairment to the standard L-dopa treatment, cognitive impairment, significant dopaminergic neurodegeneration and alpha synuclein pathophysiology. Further studies would analyze the RBD in the prodromal phase, temporal progression associating it with the neuronal degeneration as well as with the gut motility impairment to further characterize the present model. In conclusion, in comparison of the genetic models or end stage neurotoxin-based models, our chronic P+L model allows the evaluation of novel therapeutics at the early PD stages as well as for their efficacy in halting or slowing the progression of the disease.

## Acknowledgements

We thank for the additional funding was provided by the Anne M. and Philip H. Glatfelter, III Family Foundation and Ron and Pratima Gatehouse Foundation.

## Authors contribution

**Vaibhavi Peshattiwar:** Writing – original draft, Writing - Review & Editing, Visualization, Methodology, Investigation, Validation, Formal analysis, Data curation, Project administration. **Caroline Swain:** Investigation, Data Curation, Formal analysis. **Dipesh Pokharel:** Investigation, Data Curation. **Khoi Le:** Investigation, Data Curation, Formal analysis. **Isabel Kennedy:** Investigation, Data Curation. **Kala Venkiteswaran:** Writing – review & editing, Conceptualization, Methodology, Formal analysis, Data curation, Supervision, Project administration. **Thyagarajan Subramanian:** Writing – review & editing, Writing – original draft, Supervision, Resources, Project administration, Methodology, Funding acquisition, Conceptualization.

## Data availability

The data that support the findings of this study are available on request from the corresponding author.

